# Epithelial Folding Irreversibility is Controlled by Elastoplastic Transition via Mechanosensitive Actin Bracket Formation

**DOI:** 10.1101/2023.12.19.572470

**Authors:** Aki Teranishi, Misato Mori, Rihoko Ichiki, Satoshi Toda, Go Shioi, Satoru Okuda

## Abstract

During morphogenesis, epithelial sheets undergo sequential folding to form three-dimensional organ structures. The resulting folds are irreversible, ensuring that morphogenesis progresses in one direction. However, the mechanism establishing the irreversibility of folding remains unclear. Here, we report a novel mechanical property of epithelia that is responsible for folding irreversibility. Using a newly developed mechanical indentation assay, we demonstrate that short-term or low-curvature folding induces an elastic, shape-restoring response. In contrast, combined long-term, high-curvature folding results in plastic, irreversible deformation. This elastic-to-plastic transition occurs in a switch-like manner, with critical thresholds for the folding curvature and duration. Specific cells at the fold initiate this transition, sensing the curvature and duration of folding on their apical side via mechanosensitive signaling pathways, including transient receptor potential canonical (TRPC) 3/6-mediated calcium influx and ligand-independent epidermal growth factor receptor activation. These pathways induce F-actin accumulation into a bracket-like structure across the fold, establishing the transition. The duration threshold is determined and tunable by the actin polymerization rate. These results demonstrate that cells control the irreversibility of epithelial folding by detecting folding characteristics and adaptively switching between elastic and plastic responses. This finding resolves a long-standing question about the directionality of morphogenesis.

---

Morphogenesis proceeds in one direction, with sequential epithelial folds transforming planar sheets into three-dimensional structures. Importantly, each fold is irreversible, and this irreversibility is essential for the formation of complex organ structures. Morphogenesis cannot proceed if the folds return to their original shapes; therefore, ensuring the irreversibility of epithelial folding is fundamental for achieving morphogenesis.

Various cellular behaviors that generate mechanical forces, including apical constriction, serve as key drivers of epithelial folding^1–5^. The balance between these forces and mechanical properties, such as elasticity and viscosity, determines the amount and rate of tissue deformations^6–10^. However, folding irreversibility cannot be explained by either driving forces or viscoelastic properties. Because the epithelium is constantly exposed to various deformations by the activities of its constituent cells and surrounding tissues^11–13^, successful morphogenesis requires epithelia to retain necessary deformations while resisting unnecessary deformations.

We previously demonstrated that the epithelium undergoes plastic bending, facilitating retinal invagination in a mechanosensitive manner^14^. Additionally, recent studies examining tissue fluidization^15–17^ and viscoelasticity^9,18,19^ have, either implicitly or explicitly, described plastic deformations. In contrast, the epithelium exhibits bending elasticity, which allows it to resist external forces^20–22^. Cells respond mechanosensitively to imposed deformations^23–26^ and surrounding curvatures^27–29^, indicating that cells may regulate their bending elasticity and plasticity—properties that confer opposing responses to external forces—to control the irreversibility of epithelial folding. However, although the importance of mechanical plasticity is increasingly recognized^30^, the elastoplastic property of epithelial folding and its regulatory mechanisms remain poorly understood.

We developed a mechanical indentation assay for measuring the elastoplastic properties of the epithelium. We observed both elastic and plastic responses in several types of epithelia, including a mouse embryonic optic vesicle (OV), mouse embryonic stem cell (mESC)–derived OV organoid, and Madin-Darby canine kidney (MDCK) cell–derived cyst. Further experiments using the MDCK cell– derived cyst revealed an elastoplastic transition and its underlying molecular mechanisms, which likely serve to adaptively control the irreversibility of the epithelial folding response through the detection of imposed deformation.

## Epithelial response switches between elastic and plastic depending on folding duration and curvature

To measure epithelial bending plasticity, we developed an indentation assay, in which a glass pipette is used to indent the epithelium, forming a single fold. After holding the pipette in place for a defined time, the pipette is removed to relax the residual strain (**Fig. 1a, b** and **Supplementary Fig. 1a–e**). This assay enables the assessment of bending properties as a function of the hold duration (*τ*) and curvature (κ).

**Figure 1.**
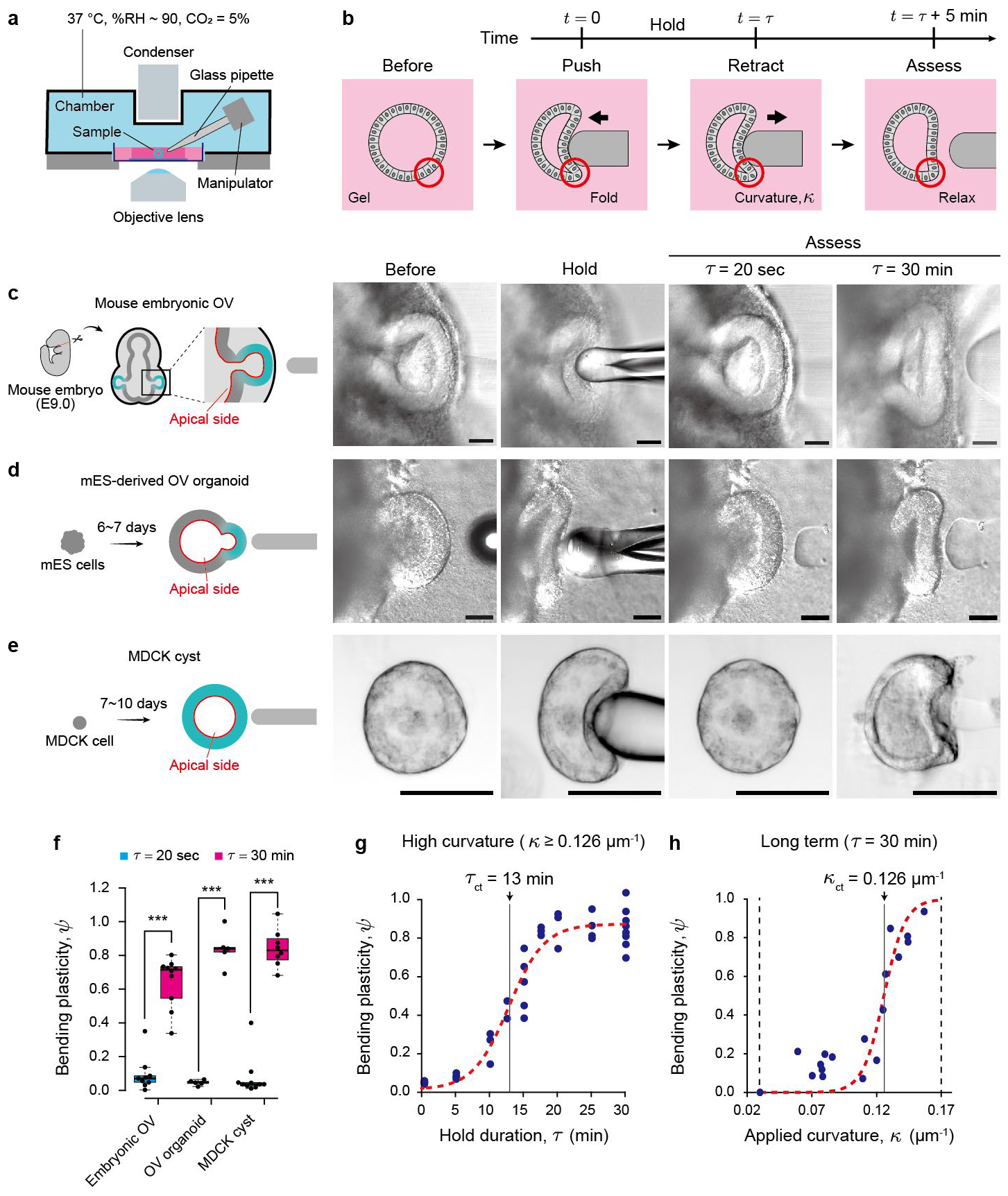
Epithelial response switches between elastic and plastic depending on folding duration and curvature. (a) Mechanical indentation device with a custom culture chamber on a confocal microscope. An electric piezo manipulator with a glass pipette is placed inside. (b) Schematic for the flow of the indentation assay. Bending plasticity, *ψ*, was introduced, and its dependence on the hold duration, *τ*, was explored. (c–e) Schematics and images of the epithelium before, during, and after indentation for (c) mouse embryonic optic vesicles (OVs) at E9.0. (d) an OV organoid derived from mouse embryonic stem cells (mESCs) at Day 6 or 7, and (e) a cyst derived from Madin-Darby canine kidney (MDCK) cells. Scale bar, 50 μm. (f) Bending plasticity, *ψ*, of three types of epithelia (N ≥ 7). Box plots in (f): center: median; bounds, 25th and 75th percentiles; whiskers, min and max; ***P < 0.001 (Welch’s t-test). (g and h) Bending plasticity, ψ, of MDCK cell–derived cysts as a function of hold duration ττ and applied curvature κ, respectively. The red dotted line shows sigmoid functions fitted by the least squares method, in which the critical duration, *τ*_ct_, is 13 min and κ_*ct*_ is 0.126 μm^−1^, respectively.

We first assessed the dependence of the bending property on *τ*. Mouse embryonic OV tissue exhibited an elastoplastic response dependent on *τ*: when *τ* = 20 sec, the tissue restored its original shape, whereas the tissue deformed irreversibly when *τ* = 30 min (**Fig. 1c**; **Video S1**). A similar response was observed in mESC–derived OV organoids (**Fig. 1d**; **Video S1**), indicating that this elastoplastic response is an intrinsic property of OVs, independent of surrounding tissues, such as the epidermis or the periocular mesenchyme. MDCK cell–derived cysts displayed a similar elastoplastic response (**Fig. 1e**; **Video S1**), suggesting that this response is not specific to mouse optic vesicles but is likely a general characteristic of many types of epithelia, including embryonic and cultured tissues. The plastic deformation that occurred after the 30-min indentation was maintained for several hours after unloading (**Fig. 1b, f**).

In experiments using *τ* = 30 min, the tissue exhibited springback, in which a slight return to the original shape was observed after retraction, indicating a mixed elastic and plastic response. To quantify this mixed response, we introduced the degree of bending plasticity, *ψ*, based on the epithelial curvature measured at the fold caused by the indentation (see **Methods**). This criterion represents the contribution of plastic deformation to total deformation. Significant differences were observed between the elastic and plastic responses among embryonic OV tissues, mESC–derived OV organoids, and MDCK cell–derived cysts (**Fig. 1f**).

We performed further quantitative analyses using MDCK cell–derived cysts, which have consistent sizes and shapes and no surrounding tissues. The measurements revealed that *ψ* is dependent on *τ* and κ, with increases in *τ* and *c* associated with a monotonic increase in *ψ*, from elastic to plastic (**Fig. 1g, h**). Remarkably, the rate of increase in *ψ* is nonlinear, with a switch-like transition observed at certain *τ* and κ values. By fitting a sigmoidal function, we identified the critical duration as *τ*_ct_ = 13 min and the critical curvature as κ_ct_ = 0.126 μm^-1^. The original shape tended to be restored when *τ* was below either *τ*_ct_ or κ_ct_, whereas the tissue deformed irreversibly when *τ* was above *τ*_ct_ and κ_ct_. These results show that the epithelium switches its bending response bistably between elastic and plastic depending on the characteristics of the imposed deformation.

## Epithelial folding triggers ligand-free activation of EGFR and downstream PI3K-Akt pathway

To understand how cells sense deformation, we measured *ψ* after the application of inhibitors. During indentation, epidermal growth factor receptor (EGFR) becomes activated in a mechanosensitive manner, and the inhibitors PD153035, LY294002, and ipatasertib, which inhibit EGFR, phosphoinositide 3-kinase (PI3K), and Akt, respectively, all suppressed the elastoplastic transition (**Fig. 2a**), indicating that the EGFR–PI3K–Akt pathway regulates the transition.

**Figure 2.**
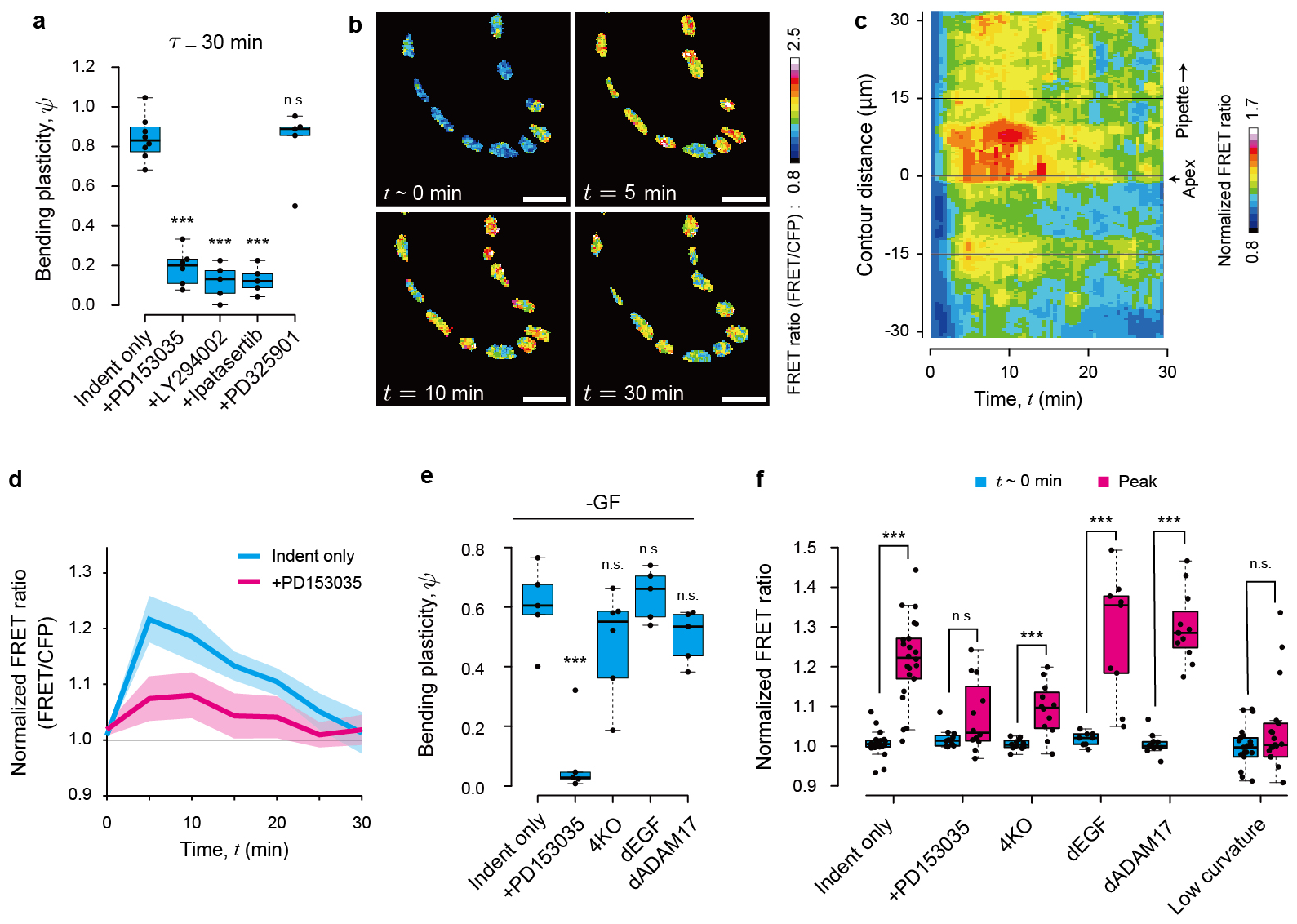
Epithelial folding triggers ligand-free activation of EGFR and downstream PI3K-Akt pathway. (a) Bending plasticity, *ψ*, for indentation only and indentation in the presence of inhibitors for EGFR or downstream molecules, at *τ* = 30 min (N ≥ 5). (b) ERK-FRET images for indentation only immediately after initiation, at the time of peak ratio, and 30 min after indentation. Scale bars, 15 μm. (c) Kymograph of ERK-FRET ratio along the epithelium contour over time under conditions of indentation only. FRET ratio at each pixel is normalized to its value immediately after indentation and averaged across samples (N = 5). The pipette is at a positive distance from the apex with the highest curvature. (d) Time variations of ERK-FRET ratios around the apex during indentation for indentation only and indentation in the presence of EGFR inhibition (N = 8). Data represent the mean ± s.e. (e) Bending plasticity, *ψ*, in cysts featuring the knockout of EGFR agonists, including all four factors (4KO), EGF (dEGF), and a factor upstream of the other three agonists (dADAM17), for *τ* = 30 min (N ≥ 5). (f) ERK-FRET ratios of cysts featuring knockout of EGFR agonists, the condition with low-curvature folding at *t* = 0 min, and the time of the peak ratio (N ≥ 5). Box plots in (a), (e), and (f): center, median; bounds, 25th and 75th percentiles; whiskers, min and max; ***P < 0.001 (Welch’s t-test). ADAM17, a disintegrin and metalloproteinase 17; EGF, epidermal growth factor; EGFR, epidermal growth factor receptor; ERK, extracellular signal-related kinase; FRET, Förster resonance energy transfer; PI3K, phosphoinositide 3-kinase.

The mitogen-activated protein kinase kinase (MEK) inhibitor PD325901 did not affect the elastoplastic transition (**Fig. 2a** and **Supplementary Fig. 2a**), indicating that the MEK–ERK pathway is not involved in the transition. However, the deformation activated extracellular signal-related kinase (ERK), and this ERK activity was inhibited by EGFR inhibitor PD153035 (**Fig. 2d, f** and **Supplementary Fig. 2a**), indicating that ERK activation occurred through EGFR. Therefore, to monitor the EGFR activation in the transition, we utilized a Förster resonance energy transfer (FRET) sensor for ERK activation (**Fig. 2b** and **Supplementary Fig. 2a**; **Video S2**). The FRET ratio increased rapidly within the first 5 min of indentation, particularly near the fold (**Fig. 2b, c**), followed by a gradual decrease (**Fig. 2d**).

To clarify what triggers EGFR activation, we assessed cysts derived from MDCK cells engineered to knock out all endogenous EGFR ligands: epidermal growth factor (EGF), transforming growth factor alpha (TGFα), heparin-binding EGF-like growth factor (HBEGF), and epiregulin (EREG)^31^. These cysts displayed an increased FRET ratio and elastoplastic transition in response to deformation (**Fig. 2e, f**), and similar results were obtained in cysts derived from MDCK cells engineered to knock out only EGF and those engineered to knock out disintegrin and metalloprotease 17 (ADAM17), in which TGFα, HBEGF, and EREG are inactivated (**Fig. 2e, f** and **Supplementary Fig. 2a-e**). Moreover, the FRET ratio was not increased by low-curvature folding (**Fig. 2f** and **Supplementary Fig. 2a, f**). These results show that cells sense the curvature of folding, and substantial folding triggers EGFR activity in a ligand-independent manner.

Our findings indicate that the elastoplastic transition involves the mechanosensitive and ligand-independent activation of the EGFR–PI3K–Akt pathway, which is consistent with previous reports that EGFR is activated without a ligand^32–34^ and in a mechanosensitive manner^35–37^.

## TRPC3/6-mediated mechanosensing triggers calcium transient at the epithelial fold under EGFR activation

Our inhibitor assays revealed that mechanosensitive calcium channels are required for elastoplastic transition. The transition was suppressed by the inhibition of mechanosensitive cation channels, including transient receptor potential canonical (TRPC)1/6, using Grammostola mechanotoxin 4 (GsMTx4)^38,39^, and by the chelation of intracellular calcium, using BAPTA-AM (**Fig. 3a** and **Supplementary Fig. 3a, b**). Specifically, the transition was suppressed by the inhibition of TRPC6 with SAR-7334; however, it still occurred even under the inhibition of TRPC1 with Pico145 (**Fig. 3a** and **Supplementary Fig. 3a, b**), indicating that TRPC6-mediated calcium influx is necessary for the transition. Moreover, TRPC3 and TRPC6 form a heteromeric complex on the apical side^40^, and inhibition of TRPC3 with Pyr3 suppressed the transition (**Fig. 3a**), suggesting that cells detect deformation via TRPC3/6-mediated calcium influx on the apical side.

**Figure 3.**
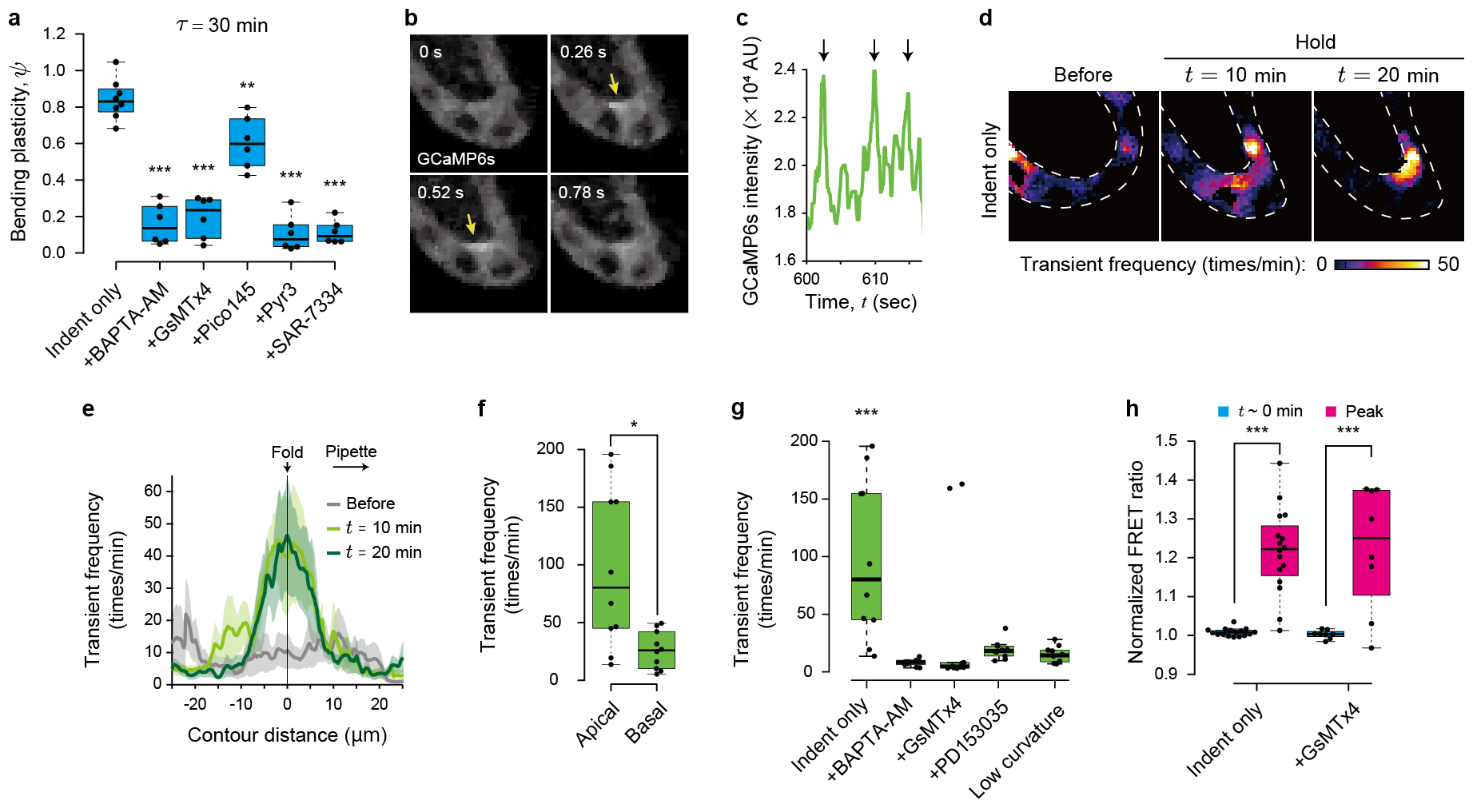
TRPC3/6-mediated mechanosensing triggers calcium transient at the epithelial fold under EGFR activation. (a) Bending plasticity, *ψ*, under conditions of mechanosensitive channel inhibition or intracellular calcium chelation for *τ* = 30 min (N ≥ 6). (b) Time-lapse images of GCaMP6s 10 min after indentation initiation. Calcium transients occurred in the apical region (arrows). (c) Time variations of calcium concentrations around the apex 10 min after indentation initiation. (d and e) Spatial distributions of calcium transient frequency: before, 10 min, and 20 min after indentation initiation. Map view (d) and distribution along the epithelium contour averaged across samples (e). Dotted lines indicate epithelium contours. Data represent the mean ± s.e. (N = 5). (f) Calcium transient frequency at apical and basal regions (N = 5). (g) Calcium transient frequency around the apex under conditions of mechanosensitive channel inhibition, intracellular calcium chelation, EGFR inhibition, or low-curvature folding (N = 5). (H) ERK-FRET ratios under conditions of mechanosensitive channel inhibition at *t* = 0 and the time of peak ratio (N = 7). Box plots in (a) and (e–h): center, median; bounds, 25th and 75th percentiles; whiskers, min and max; *P < 0.05, **P < 0.01, ***P < 0.001 (Welch’s t-test). EGFR, epidermal growth factor receptor; ERK, extracellular signal-related kinase; FRET, Förster resonance energy transfer; TRPC, transient receptor potential canonical.

We performed live intracellular calcium imaging to observe the spatial pattern of calcium influx using cysts derived from a cell line stably expressing the calcium indicator GCaMP6s. We found that indentation induced typical calcium transients (**Fig. 3b, c**; **Video S3**) that were maintained throughout the indentation duration. These transients were specifically localized at the apical side of the fold (**Fig. 3d–f**) in a region measuring approximately 20.7 μm long along the contour and approximately 5.0 μm thick along the apicobasal axis (**Supplementary Fig. 3c–e**). These transients were suppressed by GsMTx4 and BAPTA-AM (**Fig. 3g** and **Supplementary Fig. 3a, b**), which corresponds to the suppression of the transition by these inhibitors (**Fig. 3a**). Moreover, these transients were not triggered by low-curvature folding (**Fig. 3g** and **Supplementary Fig. 3a, b**). These findings show that cells detect the folding curvature on the apical side via mechanosensitive calcium influx.

We also assessed the interaction between EGFR activation and calcium influx. To clarify whether EGFR activation affects calcium signaling, we performed live calcium imaging of cysts derived from GCaMP6s-expressing cells treated with the EGFR inhibitor PD153035, which revealed that PD153035 suppressed chronic calcium transients (**Fig. 3g**, and **Supplementary Fig. 3a, b**). By contrast, we observed increased FRET in response to indentation in the presence of the mechanosensitive channel inhibitor GsMTx4 (**Fig. 3h** and **Supplementary Fig. 2**). These results indicate that the elastoplastic transition requires both mechanosensitive pathways: EGFR activation and calcium influx. While both of them are independently triggered by deformation, calcium influx is triggered under EGFR activation. Through the calcium pathway, cells detect apical deformation.

## Actin bracket formation establishes elastoplastic transition with a tunable critical duration

To elucidate the mechanism that establishes the transition, we performed live imaging of F-actin using cysts derived from an MDCK cell line, stably expressing mCherry-fused Lifeact. During indentation, F-actin gradually accumulated locally at the apical side of the fold (**Fig. 4a** and **Supplementary Fig. 4a**; **Video S4**). Three-dimensional imaging revealed F-actin accumulation not only on the apical side but also along the lateral boundaries between cells (**Fig. 4b**; **Video S5**). F-actin accumulated to form a bracket-like structure along the apical side of the epithelium, which we term the ‘actin bracket,’ approximately 9.6 μm long and 2.0 μm thick (**Fig. 4c**, d, and **Supplementary Fig. 4b-d**), corresponding to the region with frequent calcium transients (**Supplementary Fig. 3c-e**).

**Figure 4.**
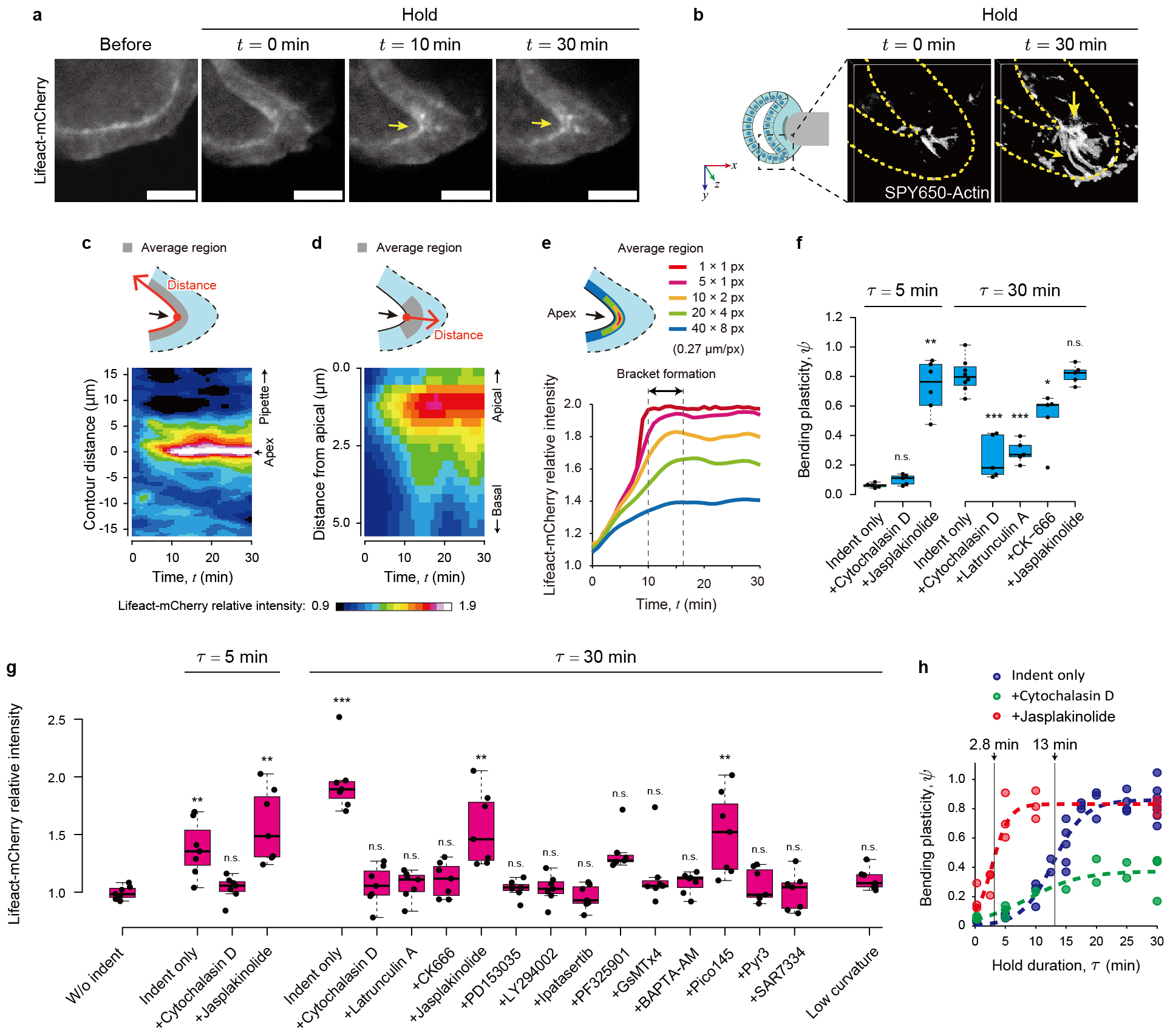
Actin bracket formation establishes elastoplastic transition with a tunable critical duration. (a) Time-lapse images of Lifeact-mCherry before indentation and at *t* = 0 min, *t* = 10 min, and *t* = 30 min. F-actin accumulated at fold (arrows). Scale bars, 15 μm. (b) Three-dimensional constructed images of SPY650-actin at *t* = 0 min and *t* = 30 min. Dotted lines indicate epithelial contours. F-actin accumulated around apical and lateral boundaries (arrows). (c and d) Kymographs of Lifeact-mCherry intensity along the epithelium contour (c) and apicobasal axis (d) over time. Intensity at each pixel was normalized to the initial intensity and averaged across samples (N = 5). (e) Time variation of the maximum and average Lifeact-mCherry intensity around fold (N = 5). (f) Bending plasticity, *ψ*, under conditions of F-actin dynamics inhibition for *τ* = 5 min and *τ* = 30 min (N ≥ 5). (g) Lifeact-mCherry intensity around the apex for *τ* = 5 min and *τ* = 30 min, under conditions of F-actin polymerization inhibition, EGFR inhibition, mechanosensitive channel inhibition, or low-curvature folding (N = 7). Values were normalized to those before indentation. (h) Bending plasticity, *ψ*, as a function of duration time, *τ*, under conditions of actin polymerization inhibition or enhancement. Dotted lines show fitted sigmoid functions. Box plots in (f and g): center, median; bounds, 25th and 75th percentiles; whiskers, min and max; *P < 0.05, **P < 0.01, ***P < 0.001 (Welch’s t-test). EGFR, epidermal growth factor receptor.

F-actin accumulation occurred most rapidly at the point of highest curvature, showing a linear increase for the first 10 min and then plateauing (**Fig. 4e**). This time duration is shorter than the critical duration for the elastoplastic transition (**Fig. 1G**). The region of accumulation gradually expanded from this point, taking approximately 16 min to fully form the actin bracket and cover the fold (**Fig. 4e**). This time duration corresponds to the critical duration of the transition (**Fig. 1g**), indicating that the formation of the actin bracket is essential for the transition.

To clarify the role of F-actin accumulation in the elastoplastic transition, we measured *ψ* following the inhibition of F-actin polymerization with cytochalasin D, latrunculin A, or CK-666 (**Fig. 4f, g and Supplementary Fig. 4e, f**). These inhibitors not only suppressed F-actin accumulation but also inhibited the transition, indicating the critical role of F-actin polymerization. In addition, inhibiting the EGFR-PI3K–Akt and calcium pathways, which are known regulators of F-actin dynamics^41–45^, also halted both F-actin accumulation and the transition (**Fig. 2a, 3a, 4g** and **Supplementary Fig. 4f**). By contrast, inhibiting MEK or TRPC1 did not affect F-actin accumulation, corresponding to the lack of effect on the transition. Moreover, this F-actin accumulation was not induced by low-curvature folding (**Fig. 4f** and **Supplementary Fig. 4f**). These results indicate that F-actin accumulation, which is regulated downstream of the EGFR–PI3K–Akt and calcium pathways triggered by substantial folding, is essential for the elastoplastic transition.

Interestingly, enhancing actin polymerization with jasplakinolide resulted in a plastic response as rapidly as 5 min after indentation (**Fig. 4f, g** and **Supplementary Fig. 4e**). Even in the presence of jasplakinolide, short-term indentations induced elastic responses, but the critical duration required for the elastoplastic transition (*τ*_ct_ = 2.8 min) was significantly shorter than under untreated conditions (**Fig. 4h**). This result indicates that the actin polymerization rate determines the critical duration required for elastoplastic transition.

These results show that F-actin accumulates across the fold, forming a bracket-like structure that establishes elastoplastic transition, and that the critical duration can be tuned by adjusting the actin polymerization rate.

## Discussion

In this study, we elucidated the mechanism underlying elastoplastic transition during epithelial folding, which occurs as an all-or-nothing response that is dependent on the curvature and duration of the imposed deformation. The transition is established by the accumulation of F-actin in response to mechanosensing pathway activation (**Supplementary Fig. 5**).

The concept of elastoplastic transition is well-known in the field of solid materials engineering. However, its mechanics in epithelial folding have not been previously reported, to our knowledge. Metals and resins typically exhibit strain-dependent elastoplastic transitions, with plastic deformation resulting from large deformations^46^. By contrast, elastoplastic transitions for fabrics and other materials are often time-dependent transitions, with the transition duration dependent on the material and environment. Transitions in these materials are generally caused by passive mechanisms, such as atomic displacement in metals, crystal exfoliation in resins, and fiber displacement in fabrics (**Supplementary Fig. 6a**). The tunable critical duration in epithelial folding, as regulated by living cells, represents a sharp contrast with the transition mechanics associated with other materials.

The elastoplastic transition of the epithelium is analogous to tissue fluidization^16,17,47,48^, as both of these processes are able to induce irreversible deformation (**Supplementary Fig. 6b**). However, fluidization requires tissue-to-phase transition from a solid to a liquid, whereas tissue remains in the solid phase during the elastoplastic transition. Kinematically, fluidization is caused by a global rearrangement of cells at the tissue scale, whereas the elastoplastic transition is caused by a change in local cell behavior and does not require global rearrangement. In the elastoplastic transition, local cell behavior alone can cause tissue-scale deformation, generating an irreversible fold structure.

The cells that drive apical constriction in epithelial folding may also undergo elastoplastic transition, responding to their own deformation (**Supplementary Fig. 6c**). This idea is supported by observations that elastoplastic transition and apical constriction share several key components, including actomyosin activity^1–3,14,49–52^, calcium signaling^14,50^, and EGFR activation^51–53^. Apical constriction occurs collectively for all cells within a specific region and period; however, individual cells are constantly generating relatively short-term forces through behaviors such as division, apoptosis, and contraction, which can introduce noise^48,54^. Elastoplastic transition may play a key role in stabilizing appropriate invaginations, filtering out the noise from short-term individual cell forces, and enabling collective cellular behaviors.

In conclusion, we demonstrated that cells actively control the irreversibility of epithelial folding through the elastoplastic transition. This observation requires the reconsideration of long-standing mechanistic concepts in development, indicating that epithelial morphogenesis proceeds not only by regulating active force generation but also by regulating deformation irreversibility to define the resulting morphology.

## Acknowledgments

We thank all the members of Okuda’s laboratory for discussions; S. Hayashi, Y. C. Wang, and F. Motegi for critical reading; M. Matsuda and K. Okamoto for valuable comments; K. Yoshida for help with experiments; and A. Matsuoka for help with image analysis. This work was supported by the WISE Program for Nano-Precision Medicine, Science, and Technology of Kanazawa University, Ministry of Education, Culture, Sports, Science and Technology (MEXT; to AT); the Japan Science and Technology Agency (JST), CREST [Grant No. JPMJCR1921]; the Japan Agency for Medical Research and Development [AMED; Grant No. 21bm0704065h0003]; the Japan Society for the Promotion of Science (JSPS), KAKENHI [Grants No. 21H01209, 21KK0134, 22K18749, and 22H05170]; and the World Premier International Research Center Initiative, MEXT, Japan (to SO).

## Contributions

A.T.: data curation, formal analysis, investigation, methodology, resources, software, validation, visualization, and writing (original draft and revised draft). M.M.: formal analysis and investigation. R.I.: investigation and validation. S.T.: resources and validation. G.S.: resources and validation. S.O.: conceptualization, data curation, formal analysis, funding acquisition, methodology, project administration, resources, software, supervision, visualization, and writing (original draft and revised draft).

## Methods

### Mice

All mouse studies were performed with approval from the Ethics Committee at Kanazawa University. We used Slc:ICR mice purchased from Sankyo Labo Service Co. (Toyama, Japan). We defined the midnight preceding the detection of a plug as embryonic day 0.0 (E0.0), and all mice were euthanized by cervical dislocation.

### Cell culture

Madin-Darby canine kidney (MDCK) cells were obtained from the Mostov Lab at the University of California, San Francisco. Cell lines stably expressing Lifeact-mCherry and GCaMP6s stable were established. MDCK-WT-EKARrEV, MDCK-4KO-EKARrEV, MDCK-dADAM17-EKARrEV, and MDCK-dEGF-EKARrEV cells expressing a FRET biosensor for ERK activity were contributed by Dr. Michiyuki Matsuda (Kyoto University, Kyoto)^31^. These cells were maintained in Dulbecco’s Modified Eagle Medium (DMEM, Wako) containing 10% FBS (Gibco or NICHIREI) and 1% penicillin-streptomycin (Wako) in a 5% CO_2_ incubator at 37°C.

Mouse embryonic stem cells (mESCs, EB5, Rx-GFP) were maintained as described in previous studies^55,56^. We used a subline of the mESC line EB5 (129/Ola), in which the GFP gene was knocked in under the Rx promoter (RIKEN BioResource Research Center, ID: AES0145).

### Establishment of stable cell lines

To establish MDCK cells that stably express Lifeact-mCherry, a lentiviral transduction system was used. The lentivirus was produced by co-transfection of the expression vector coding Lifeact-mCherry and lentiviral packaging plasmids (pCMVdR8.91 and pMD2.G) using PEI MAX (Polysciences #24765) into HEK293T cells. The viral supernatant was collected 48 hr after the co-transfection and filtered through a 0.45 μm filter. Then MDCK cells were infected with the viral supernatant in the presence of hexadimethrine bromide (Sigma-Aldrich #H9268). After a few days, fluorescent cells were sorted using a cell sorter (FACSMelodyTM, BD Bioscience).

To establish MDCK cells that stably express GCaMP6s, a PiggyBac transposase system was used. The expression vector coding GCaMP6s was introduced into MDCK cells with Super PiggyBac transposase (System Bioscience). After a few days, fluorescent cells were sorted using a cell sorter (SH800, SONY).

### Plasmids

Lifeact-mCherry^57^ was obtained from Michael Davidson (Addgene plasmid #54491). The coding sequence of Lifeact-mCherry was amplified with the primers 5’-AATTCTCACGCGGCCGCCACCATGGGCGTGGCCGACTTG-3’ and 5’-TCAAGCTTGCATGCCTGCAGGTTACTTGTACAG CTCGTCCATGCCGC-3’, and cloned via In-Fusion HD cloning (Clontech #Z9648N) into a modified pHR’SIN:CSW vector under an EF1α promoter.

The GCaMP6s coding sequence amplified from pGP-CMV-GCaMP6s^57^ purchased from Addgene (#40753) with the primers 5’-GGGGAATTCGCCACCATGGGTTCTCA-3’ and 5’-CCC GCGGCCGCTCACTTCGCTGTCATCA-3’ was subcloned into EcoRI and NotI sites in the P B-CMV-MCS-EF1α-Puro PiggyBac transposon vector (PB510B-1, System Bioscience).

### Three-dimensional culture for MDCK cell–derived cyst generation

Epithelial cysts were derived from MDCK cells according to the previously reported method^58,59^. Matrigel (50 μl, Corning) was polymerized on a 15-mm diameter cover glass (MATSUNAMI) at the bottom of a 24-well plate. Cell suspension containing 2% Matrigel was poured into the culture medium to obtain a final cell density of 10,000 cells/cm^2^. After approximately 7 days of culture, spherical epithelial cysts with a lumen were obtained in the gel on the cover glass.

### SFEBq for optic vesicle organoid generation

The SFEBq culture method was performed as described in a previous study^55^. In this culture, 3,000 suspended cells and 3.5% Matrigel (Corning) were added to each well of a 96-well plate to form cell aggregates. Tissues from Day 6 or Day 7, with protruding Rx-positive regions, were used for the indentation assay.

### Mouse embryonic OV culture

Mice were dissected at E9.0, and the heads were cut off and cultured in DMEM Ham’s/F’12 (Wako) supplemented with 1% N2 (Gibco) and 1% penicillin-streptomycin (Wako) in a 5% CO_2_ incubator at 37°C, as described in a previous study^14^. For the mechanical indentation assay, heads were used within 6 h of the start of incubation.

### Mechanical indentation assay

To perform the mechanical indentation assay under well-controlled conditions, we developed a custom incubator placed on the stage of a confocal microscope (LSM800, Zeiss). This incubator was maintained at conditions of 5% CO_2_, 37°C, and ≥90% humidity. In addition, we installed a three-axis piezoelectric manipulator (MM3A-LA, Kleindiek Nanotechnik) equipped with a pipette inside the incubator. To avoid adhesion of the pipette to tissues, the tip surface was coated with methacryloyloxyethyl phosphorylcholine polymers (NovyCoat, FastGene).

For indentation assays using mouse embryonic OV tissue and mESC-derived OV organoids, samples were transferred to 35-mm diameter glass-bottom dishes and embedded in Matrigel with 1 ml of culture medium. For indentation assays using MDCK cell–derived cysts, single-layer, single-lumen cysts with an average diameter (55 ± 6 μm) were selected from among 7–10-day-old cysts. The cysts were cultured in Matrigel on a coverslip, which was transferred to a 35-mm-diameter dish filled with 1 ml culture medium. Using the manipulator, a pipette was used to indent the epithelium to induce a fold at t = 0, held for the duration of *τ*, and retracted at *t* = *τ*. The epithelium was indented to induce a fold with a high amount of curvature, with the outer curvature κ greater than 0.125 μm^-1^. However, when examining the effect of the curvature of folding, the epithelium was indented to induce a lower amount of curvature, with κ less than 0.10 μm^-1^. After retraction, the epithelium reached a stable shape within 5 min. Therefore, the epithelium shape was evaluated at *t* = *τ* + 5 min. For *τ* = 30 min, the stable shape was maintained for 2–3 h after retraction. Thereafter, the shape of the epithelium gradually changed due to cell migration.

For FRET and calcium imaging, cysts were resuspended in a collagen gel to eliminate growth factors and facilitate the detection of sensitive signals. Cysts were incubated with cell recovery solution (Corning) on ice for 20 min to depolymerize Matrigel and then collected by centrifugation at 300 × *g* for 3 min. Cysts were suspended in 50 μl collagen solution consisting of 70% collagen (0.3% CellMatrix Type I-A, Nitta Gelatin), 20% Hank’s Balanced Salt Solution (HBSS, without phenol red, Wako), and 10% reconstitution buffer (Nitta Gelatin). Cysts in collagen solution were placed in a 35-mm-diameter glass-bottom dish, and collagen was allowed to polymerize at 37°C for 20 min. The dish was filled with 1 ml culture medium.

### Quantification of curvature and intensity

The curvature and fluorescence intensity along the epithelium contour were calculated as described in previous studies^60,61^. For this calculation, a mask image of the epithelium contour was manually obtained from the transmitted light image of the tissue. The curvature was calculated using the sequential three-point method. Fluorescence intensity was calculated as the average in the direction normal to the contour.

### Quantification of epithelial bending plasticity

To quantify the mechanical response of the epithelium, we introduced a parameter, *ψ*, to represent the degree of bending plasticity. This parameter is determined using the maximum curvature on the epithelium contour, specifically at a location we refer to as the apex. We measured curvature at the apex at *t* = *τ* and *t* = *τ* + 5 min. We refer to the curvature at *t* = *τ* as the curvature of the applied fold, κ, whereas the curvature at *t* = *τ* + 5 min is the curvature of the relaxed shape, κ_1_. In addition, we identified the pre-indentation location on the epithelium that corresponds to the apex immediately after the indentation begins (*t* = 0). We measured the average curvature around this location, represented by κ_0_. The degree of bending plasticity, *ψ*, is defined as

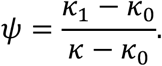

The value for bending plasticity ranges from 0–1. When *ψ* = 0, the epithelium responds completely elastically, retaining its original shape after pipette removal. Conversely, when *ψ* = 1, the epithelium responds completely plastically, maintaining the applied fold.

### Sigmoid curve fitting to bending plasticity

We employed sigmoidal functions to evaluate the dependence of bending plasticity on τ and κ. The functions fitted to the dependences on τ and κ, represented by ψ_τ_ and ψ_κ_, are described by

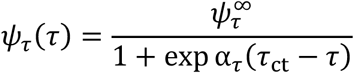

and

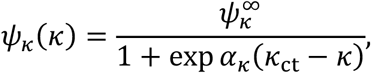

respectively. Here, 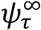, α_τ_, τ_ct_, 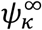, α_κ_, and κ_ct_ are constants, and τ_ct_ and κ_ct_ represent the critical duration and curvature, respectively. By fitting these equations to the data using the least squares method, we determined the values of τ_ct_ and κ_ct_.

### Three-dimensional imaging of F-actin

Three-dimensional imaging was performed to observe changes in the three-dimensional structure of accumulated F-actin at the fold during the indentation. Cysts were incubated in a culture medium containing SPY650-actin (Cytoskeleton, Inc., 1:2000 dilution) for 3 h before indentation. Fluorescence observations were made by confocal microscopy immediately after pipette indention and during holding.

### Pharmaceutical interventions

To examine molecular activities, cysts were treated with the actin polymerization inhibitors cytochalasin D (Abcam, 10 μM, 60-min incubation) or latrunculin A (Wako, 10 μM, 60-min incubation); the ARP2/3 complex inhibitor CK666 (Merck, 100 μM, 60-min incubation); the actin polymerization inducer and actin depolymerization inhibitor jasplakinolide (Abcam, 1 μM, 60-min incubation); the EGFR inhibitor PD153035 (TCI, 1 μM, overnight [O/N] incubation); the MEK inhibitor PF325901 (MedChemExpress, 10 μM, 60-min incubation); the PI3K inhibitor LY294002 (MedChemExpress, 50 μM, O/N incubation); the Akt inhibitor ipatasertib (MedChemExpress, 1 μM, O/N incubation); the mechanosensitive cation channel inhibitor (e.g., Piezo1, TRPC1/6) GsMTx4 (Abcam or Peptide Institute, 2.5 μM, 15-min incubation)l the intracellular calcium chelator BAPTA-AM (Tronto Research Chemicals, 25 μM, 60-min incubation); the TRPC1 inhibitor Pico145 (Adooq Bioscience, 1 nM, 15-min incubation); the TRPC3 inhibitor Pyr3 (Merck, 3 μM, 15-min incubation); or the TRPC6 inhibitor SAR-7334 (Selleck, 100 nM, 15-min incubation).

### Confocal FRET imaging

FRET images were obtained and processed using the method described in our previous study^62^. To eliminate growth factors from the gel and enable the detection of sensitive signals, cysts were resuspended in a collagen gel before performing FRET imaging, and media was replaced with HBSS supplemented with calcium and magnesium and without phenol red at least 1 h before the indentation assay to eliminate growth factors from the media. Time-lapse images were acquired using a confocal microscope (FV-3000, Olympus) equipped with a 20× Objective lens (20×-UPlanXApo, Olympus), a GaAsP detector (FV31-HSD, Evident), and a dichroic mirror (U-FBNA, Olympus). A 405 nm laser was used to excite cyan fluorescent protein (CFP). Emission was detected at 460–480 nm for CFP (donor) and 510–530 nm for yellow fluorescent protein (YFP, acceptor). The FRET/CFP ratio was calculated after subtracting the background, and only regions with high YFP signal were visualized using ImageJ.

### Calcium transient measurement

To observe calcium transients, cysts stably expressing GCaMP6s were resuspended in collagen gel. Time-lapse images were acquired for 1 min at each of three time points: before, 10 min after, and 20 min after indentation. To detect fast calcium responses, acquisition conditions were set to 128 × 128 pixels, with a frame interval of approximately 130 ms.

To quantify the calcium transient frequency, GCaMP6s images were processed using ImageJ. Prior to processing, each image was smoothed by calculating the median value over each frame and time period at each pixel. The base fluorescence intensity level at each pixel was subtracted from the images and calculated as a moving average in the time direction over a width of 50 frames before and 50 frames after the central image. The number of calcium transients was calculated as the number of times that the difference between adjacent time frames at each pixel exceeded a value 15% higher than the average of all time frame images.

The distribution of calcium transient frequency along the epithelial contour over time was determined by normalizing the value of each pixel against the ratio of the average of each experiment to the average of all experiments, averaging the frequency in 5 pixels from the apical side in each experiment, and calculating statistical values between multiple experiments. Statistical values comparing indentation alone with indentation in the presence of pharmaceutical inhibitors and comparing the apical and basal sides were calculated by normalizing the value of each pixel against the ratio of the average before to the average after indentation for each experiment. The data after indentation includes measurements conducted at 10 min and 20 min after indentation. The length and thickness of the region in which a high frequency of calcium transients was detected were estimated by fitting Gaussian functions to the frequency distributions along the contour and apicobasal axes, respectively, centered on the maximum value in the average map.

### Statistical analysis of F-actin distribution

To quantify the spatial distribution of F-actin, Lifeact-mCherry images were processed using ImageJ. Each image was smoothed by calculating the median value over each frame and time period at each pixel. The epithelial contour was manually determined, and each fluorescence image was transformed into a map along the contour and apicobasal axes. The resulting maps were averaged across experiments by aligning each apex with the highest curvature along the contour axis and each apical surface along the apicobasal axis. The average value around the apex was calculated from the average map by defining regions of 5 × 1, 10 × 2, 20 × 4, and 40 × 8 pixels. The regions were set around the pixel with the highest intensity at the apex. The length and thickness of each actin bracket were estimated by fitting Gaussian functions to the intensity distributions along the contour and apicobasal axes, respectively, centered on the maximum value in the average map.

## Data availability

Data collected and computer codes developed for this study are available on request to the corresponding author.

## Supplemental information

**Video S1** Mechanical indentation assay and elastoplastic response.

**Video S2** Live imaging of EGFR activity with ERK-FRET sensor during indentation.

**Video S3** Live imaging of calcium dynamics with GCaMP6s during indentation.

**Video S4** Live imaging of F-actin dynamics with Lifeact-mCherry during indentation.

**Video S5** Three-dimensional structure of accumulated F-actin at the fold.

